# A Reversible Mitochondrial ROS Probe for Monitoring Mitophagy Dynamics: Development and Application of MitoFlare

**DOI:** 10.1101/2025.11.18.689025

**Authors:** Shanshan Hou, Xin Yan, Xiaoxu Li, Jingyue Ju, Zhiying Shan, Lanrong Bi

**Affiliations:** Department of Chemistry, Michigan Technological University, Houghton, Michigan 49931, United States; Department of Kinesiology and Integrative Physiology, Michigan Technological University, Houghton, Michigan 49931, United States; Health Research Institute, Michigan Technological University, Houghton, Michigan 49931, United States; Department of Molecular Pharmacology and Therapeutics and Department of Chemical Engineering, Columbia University, New York City, NY 10027, United States

**Author notes:** Corresponding Authors: Jingyue Ju, Zhiying Shan and Lanrong Bi.

**Keywords:** Mitochondrial oxidative stress, mitophagy dynamics, reversible mitochondrial ROS probe

## Abstract

Mitochondrial dysfunction and defective mitophagy are defining features of numerous neurodegenerative and metabolic disorders, yet existing tools provide limited ability to quantify mitophagy dynamics in real time within living, post-mitotic cells. Here we present MitoFlare, a mitochondria-targeted, reversible mtROS-responsive fluorogenic probe that enables continuous, non-genetic visualization of mitochondrial oxidative activation and turnover. MitoFlare incorporates dual TEMPO nitroxide quenchers into a long-wavelength rhodamine scaffold, producing >95% basal quenching and rapid, fully reversible fluorescence activation in response to mitochondrial superoxide, hydroxyl radicals, lipid-derived peroxyl species, and peroxynitrite. When combined with LysoTracker Green, MitoFlare forms a dual-probe imaging platform that resolves the entire mitophagy cascade with high spatial and temporal fidelity in intact PC12 neuronal cells.

Using this platform, we established a quantitative framework comprising three mechanistically distinct metrics: (i) a proximity index that reports early mitochondrial engagement with lysosomes, (ii) Manders’ M1 coefficient that captures mid-stage mitochondria–lysosome fusion and mitophagosome formation, and (iii) a quenching/swelling index that resolves terminal lysosomal degradation. Nutrient deprivation induced a complete, temporally ordered mitophagy program, including mtROS priming, Parkin–OPTN-associated fusion, and efficient acidification-dependent cargo degradation. In contrast, inhibition of v-ATPase with bafilomycin A1 arrested mitophagy at the fusion stage, resulting in persistent redox-active mitochondrial cargo that failed to undergo lysosomal digestion. Importantly, MitoFlare’s reversible redox chemistry uniquely revealed accumulation of undegraded, oxidatively active mitochondrial remnants within non-acidified vesicles—pathological intermediates that are undetectable using irreversible ROS dyes or genetically encoded reporters.

These findings demonstrate that mitophagy proceeds through discrete, redox-regulated and lysosome-dependent phases that can be quantitatively mapped in real time. By enabling synchronized measurement of oxidative activation, organelle trafficking, fusion, and degradation, the MitoFlare–LysoTracker system establishes a new benchmark for dynamic mitophagy analysis in physiologically relevant models. This platform provides a powerful foundation for mechanistic interrogation of mitochondrial quality control and for accelerating the discovery of therapeutic strategies aimed at restoring mitophagic fidelity in neurodegenerative, cardiovascular, and metabolic diseases.

## Introduction

Mitochondrial dysfunction is a central driver of human disease and mortality and represents a growing global health burden that urgently demands improved diagnostic and therapeutic strategies. In energy-intensive organs such as the brain, heart, and liver, mitochondria play indispensable roles in ATP production, calcium buffering, and metabolic signaling. Even brief disruptions in mitochondrial quality control can result in bioenergetic collapse, oxidative damage, and cell death. Among the multiple mechanisms preserving mitochondrial integrity, mitophagy—the selective autophagic removal of damaged mitochondria—is crucial for maintaining redox balance and cellular homeostasis. Impairment of this process has been implicated in numerous disorders, including Parkinson’s disease (PD), Alzheimer’s disease (AD), and amyotrophic lateral sclerosis (ALS), where insufficient mitochondrial clearance promotes protein aggregation, neuroinflammation, and synaptic failure.^1–6^ With neurodegenerative disease prevalence projected to rise sharply by 2050, elucidating how mitophagy is spatially and temporally regulated, particularly at its redox-controlled initiation phase, has become an urgent priority.^7–11^ However, efforts to define mitophagic dynamics in intact systems such as primary neurons, iPSC-derived cardiomyocytes, and ex vivo brain slices remain limited by the lack of reliable tools for non-genetic, real-time tracking of mitochondrial turnover.

Mitochondrial reactive oxygen species (mtROS) are not merely byproducts of metabolism but act as key regulatory signals that initiate mitophagy. Species such as superoxide (O₂^-^), hydrogen peroxide (H₂O₂), peroxynitrite (ONOO^-^), hydroxyl radicals (•OH), and lipid peroxyl radicals (ROO•) serve dual roles—reflecting mitochondrial stress while triggering selective clearance pathways.^7–11^ mtROS accumulation promotes the stabilization of PINK1 on the outer mitochondrial membrane, which recruits and activates the E3 ubiquitin ligase Parkin, leading to the ubiquitination of mitochondrial surface proteins and subsequent engagement with LC3-positive autophagosomes.^12–16^ In parallel, PINK1–Parkin-independent mechanisms such as BNIP3- and NIX-mediated signaling promote mitophagy under hypoxia or nutrient stress by directly tethering mitochondria to autophagic membranes.^17,18^ The interplay of these pathways occurs on rapid time scales and within discrete subcellular domains, underscoring the need for imaging tools capable of simultaneously capturing mtROS generation and mitochondrial degradation in living cells.

Existing mitophagy detection methods face substantial limitations. Common genetic reporters such as GFP-LC3 or tandem fluorescent LC3 variants provide only indirect readouts of autophagy progression and lack mitochondrial specificity.^19^ Their reliance on transfection or viral expression restricts their utility in post-mitotic systems, where gene delivery efficiency is low and expression variability confounds quantification. Furthermore, these reporters require separate mitochondrial and lysosomal dyes, increasing the likelihood of spectral crosstalk and phototoxicity during prolonged imaging sessions. Conventional mtROS probes such as MitoSOX Red, while useful for detecting superoxide, are irreversibly oxidized, exhibit off-target reactivity, and cannot distinguish reversible redox signaling from terminal oxidative injury.^20,21^ Consequently, the field has lacked a method that directly links real-time mitochondrial redox events to subsequent mitophagic clearance.

To address these limitations, we developed MitoFlare, a mitochondria-targeted, reversible fluorogenic probe designed for non-genetic, live-cell monitoring of mitophagy dynamics. MitoFlare incorporates dual TEMPO (2,2,6,6-tetramethylpiperidine-1-oxyl) nitroxide quenchers conjugated to a rhodamine core. In healthy mitochondria, fluorescence is efficiently quenched through spin exchange; upon exposure to mtROS species such as O₂^-^, •OH, ROO•, or ONOO^-^, the nitroxides are reduced to hydroxylamines, restoring the rhodamine’s fluorescence.^22,23^ This reversible redox chemistry enables repeated cycles of fluorescence activation and quenching, allowing MitoFlare to report dynamic changes in mitochondrial oxidative state and recovery—capabilities unattainable with conventional irreversible dyes.

The rhodamine scaffold of MitoFlare provides exceptional brightness, low autofluorescence, and minimal phototoxicity, enabling extended live-cell imaging over hours without perturbing mitochondrial physiology.^22,23^ When combined with LysoTracker, which labels acidic lysosomal compartments, the dual-probe system provides a direct window into mitophagy progression—from the onset of mtROS elevation to lysosomal fusion and degradation. Ratiometric detection further enhances reproducibility by compensating for variations in probe loading and illumination intensity.^24^ In contrast to previous tools, MitoFlare uniquely bridges the redox and degradative dimensions of mitophagy. Its reversible, ROS-specific activation captures the temporal sequence of mitochondrial stress, recruitment, fusion, and degradation, while its spectral compatibility with LysoTracker enables simultaneous quantification of oxidative signaling and lysosomal processing within the same live-cell experiment. This integration allows for precise measurement of mitophagy flux kinetics under physiological and pathological conditions, including early mitochondrial priming, mid-stage autophagosome fusion, and late-stage lysosomal clearance. MitoFlare represents a conceptual and technical advance in the study of mitochondrial quality control. By providing a non-genetic, ratiometric, and reversible means to visualize mtROS-regulated mitophagy in real time, it offers a powerful platform to dissect the redox checkpoints that govern mitochondrial turnover.

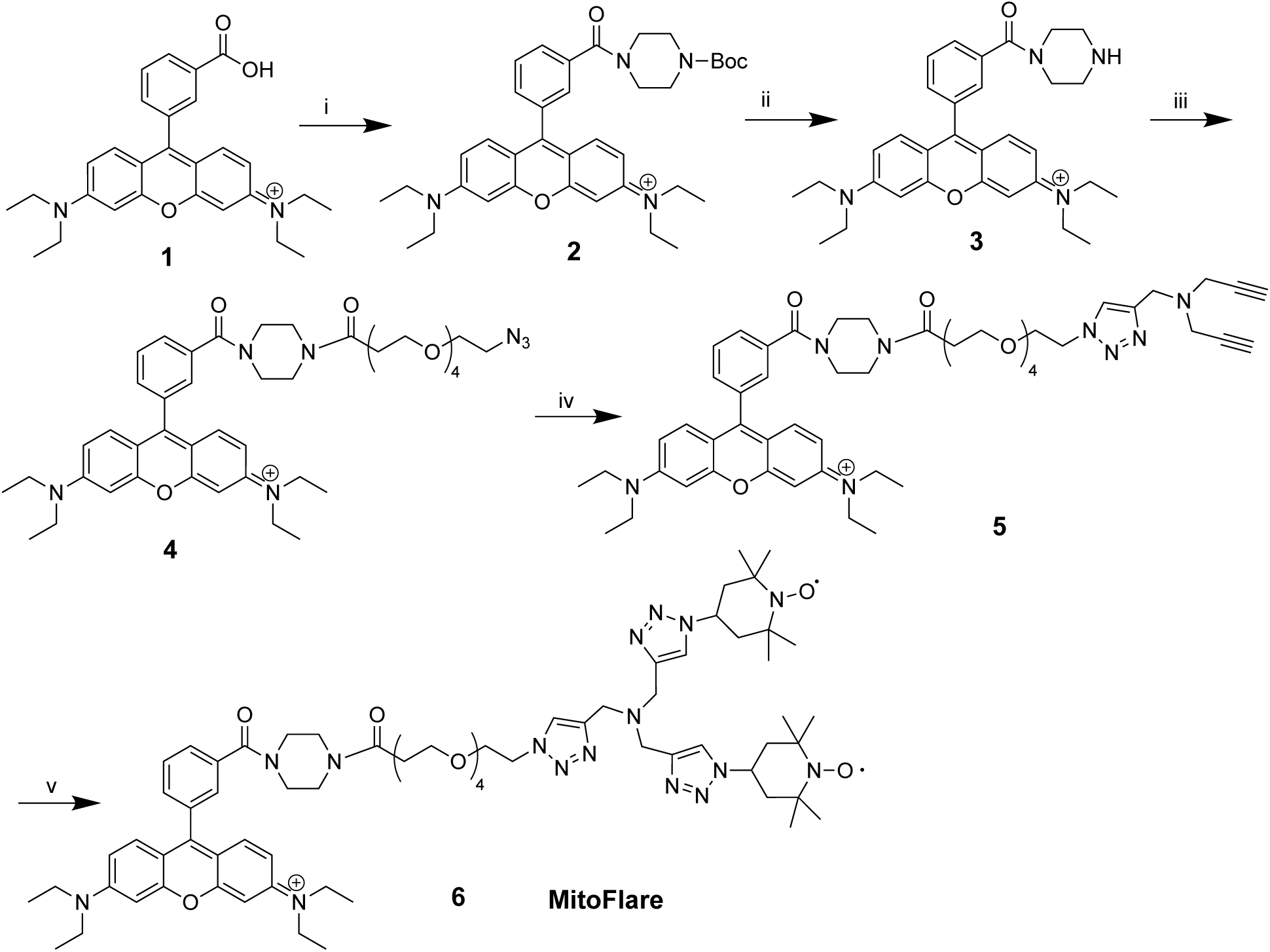

## 2. Results and Discussion

### 2.1. Chemical Design Rationale

MitoFlare was conceived as a modular fluorescent probe that combines reversible mtROS sensing, efficient mitochondrial targeting, and photostable red fluorescence to support real-time tracking of mitophagy with high fidelity. Its design integrates three functional elements—a redox-responsive sensing domain, a mitochondrial localization framework, and a rhodamine-based reporter core—each selected to provide robust performance in live-cell imaging environments where mitochondrial turnover unfolds over minutes to hours.

The redox-sensing module consists of two TEMPO (2,2,6,6-tetramethylpiperidine-1-oxyl) nitroxide moieties, introduced onto the fluorophore through copper-catalyzed azide–alkyne cycloaddition (CuAAC). These nitroxide radicals react selectively with several physiologically relevant mtROS species—including superoxide, peroxynitrite, hydroxyl radicals, and lipid peroxyl radicals—undergoing one-electron oxidation or reduction to form non-paramagnetic products.^22,23^ In the basal state, the paired TEMPO groups efficiently quench rhodamine fluorescence by photoinduced electron transfer (PET), suppressing background signal and maintaining the probe in a dark “off” configuration. Upon encountering ROS, this quenching pathway is disrupted and fluorescence is restored. ^22,23^ The dual-radical arrangement strengthens both basal quenching and sensitivity, producing a broad dynamic range and allowing the probe to detect subtle shifts in mitochondrial redox state that accompany early mitophagic priming. Importantly, this process is reversible, allowing the same mitochondrial population to be monitored through successive cycles of oxidative activation and recovery—an essential feature for studying mitophagy kinetics.

MitoFlare localizes to mitochondria through the physicochemical properties of its rhodamine scaffold. Rhodamine derivatives accumulate electrophoretically across the mitochondrial inner membrane due to their delocalized positive charge and lipophilicity, achieving substantial intramitochondrial enrichment at physiological membrane potentials. The appended TEMPO groups further increase hydrophobicity, assisting membrane passage, while a flexible tetraethylene glycol (PEG4) linker enhances aqueous solubility and minimizes aggregation. This architecture positions the redox-reactive nitroxides within the mitochondrial matrix, ensuring that fluorescence activation directly reflects local oxidative events associated with mitophagy initiation.

The rhodamine core was chosen for its favorable optical and photophysical properties in live-cell imaging. Excitation and emission in the ∼560/590 nm range reduce background autofluorescence and minimize phototoxicity relative to shorter-wavelength probes, enabling extended time-lapse imaging in delicate preparations. High brightness, strong photostability, and well-defined spectral separation support reliable quantification during multi-hour imaging sessions. In addition, ratiometric behavior arising from changes in emission characteristics during dequenching provides a built-in correction for probe loading variability and illumination differences, improving quantitative accuracy in flux measurements.

Synthesis follows a streamlined modular route in which a bis-alkyne PEG4–rhodamine precursor (compound 5, Scheme 1) undergoes sequential CuAAC reactions with azido-TEMPO reagents. This strategy affords high yields and consistent product formation while enforcing the rigid triazole geometry needed for efficient PET-based quenching. The PEG4 spacer maintains an optimal distance between the fluorophore and nitroxides, balancing quenching efficacy with conformational flexibility to preserve mitochondrial uptake.

This chemical blueprint directly addresses the shortcomings of commonly used mitochondrial ROS indicators, which often rely on irreversible redox transformations, exhibit narrow specificity, or suffer from photobleaching and off-target reactivity. By integrating reversible TEMPO-based sensing with the targeting and optical strengths of the rhodamine platform, MitoFlare provides a reliable and quantitatively precise tool for monitoring mitochondrial redox signaling and its coupling to mitophagy progression in real time.

### 2.2. Solution-Phase Assessment of MitoFlare Reversibility by Spectrofluorometric Cycling

To validate the reversible redox sensing of MitoFlare, we conducted redox cycling assay mimicking transient mtROS bursts during mitophagy. Time-course fluorescence measurements of 10 μM MitoFlare in PBS (pH 7.4) revealed robust, iterative activation across three cycles of H₂O₂ induction and ascorbic acid quenching. Each cycle elicited a ∼40-fold fluorescence surge, plateauing within 5–10 min post-induction, followed by 85–90% quenching efficiency upon antioxidant addition. Cumulative signal attenuation remained minimal (<5% over 60 min), with peak fold-changes showing no significant variance. In parallel, a probe-alone control exhibited <2% baseline drift, confirming photostability (Figure 1).

**Figure 1.**
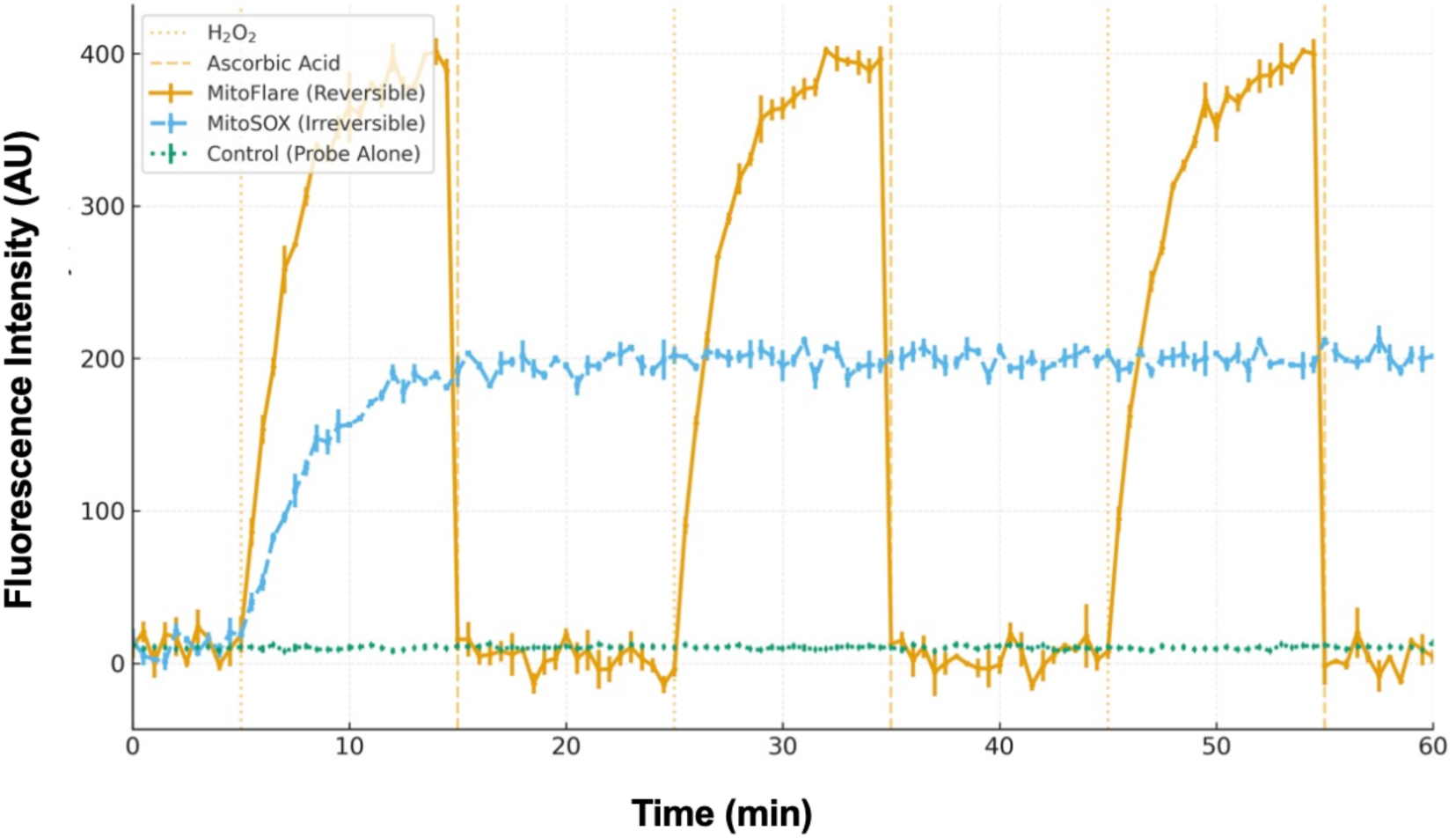
Solution-phase cycling assay demonstrates rapid, reversible redox switching of MitoFlare compared with irreversible oxidation of MitoSOX. Fluorescence time-course traces (mean ± SEM, n = 3) of MitoFlare (10 μM)/MitoSOX (10 μM), and MitoFlare-alone controls recorded in aqueous solution under three sequential redox cycles. Oxidative activation was induced by addition of H₂O₂ (100 μM at 5, 25, and 45 min), followed by reductive quenching with ascorbic acid (1 mM at 15, 35, and 55 min). MitoFlare exhibited rapid, ∼40-fold exponential fluorescence activation during each H₂O₂ pulse and robust quenching (∼85–90% signal reversal) upon ascorbate addition, with nearly identical amplitudes across cycles, indicating minimal photochemical or chemical fatigue. In contrast, MitoSOX showed a monotonic, cumulative fluorescence increase that did not respond to reductive quenching, consistent with its irreversible oxidation mechanism. Probe alone displayed minimal drift, confirming assay stability. These data establish MitoFlare as a chemically reversible, repeatedly resettable ROS reporter, in stark contrast to traditional irreversible probes, enabling precise quantification of oscillatory redox events and dynamic mitochondrial ROS flux.

These data affirm MitoFlare’s TEMPO-mediated reversibility, where dual nitroxide quenching via PET yields a high dynamic range without irreversible commitment, surpassing one-shot indicators like MitoSOX Red. The sustained cycling fidelity—rooted in hydroxylamine reoxidation—enables quantitative tracking of mtROS oscillations, critical for dissecting PINK1/Parkin flux in live models. This non-genetic tool thus bridges redox priming and autophagic resolution, offering translational potential for mitophagy-targeted therapies in oxidative pathologies.

### 2.3. Cellular Assessment of MitoFlare Reversibility in Live PC12 Neuronal Cells

To determine whether MitoFlare’s reversible redox behavior is preserved in a physiological setting, we evaluated its performance in a live-cell ROS induction–quenching assay using differentiated PC12 neuronal cells, a model that reliably recapitulates mitochondrial stress relevant to neurodegenerative pathology. Cells preloaded with 1 μM MitoFlare were exposed to three sequential rounds of antimycin A–induced mitochondrial ROS generation (5 μM at 10, 55, and 80 min), each followed by reductive quenching with N-acetylcysteine (NAC, 0.5 mM at 35, 80, and 105 min). Real-time fluorimetry revealed pronounced and reproducible cycling of MitoFlare fluorescence throughout the experiment (Figure 2). Each antimycin pulse elicited a rapid ∼2.4-fold activation, reaching maximal fluorescence within ∼30 min, consistent with robust superoxide surges triggered by Complex III inhibition. NAC addition produced a corresponding 60–70% signal decline over ∼20 min, demonstrating efficient and highly reproducible probe re-reduction. Across all three cycles, fluorescence amplitudes remained stable, with <8% cumulative attenuation over 120 min (n = 6 wells) and no significant differences in peak values (one-way ANOVA, F = 1.12, P = 0.35), confirming that MitoFlare retains full redox reversibility during repeated mitochondrial stress–recovery transitions.

**Figure 2.**
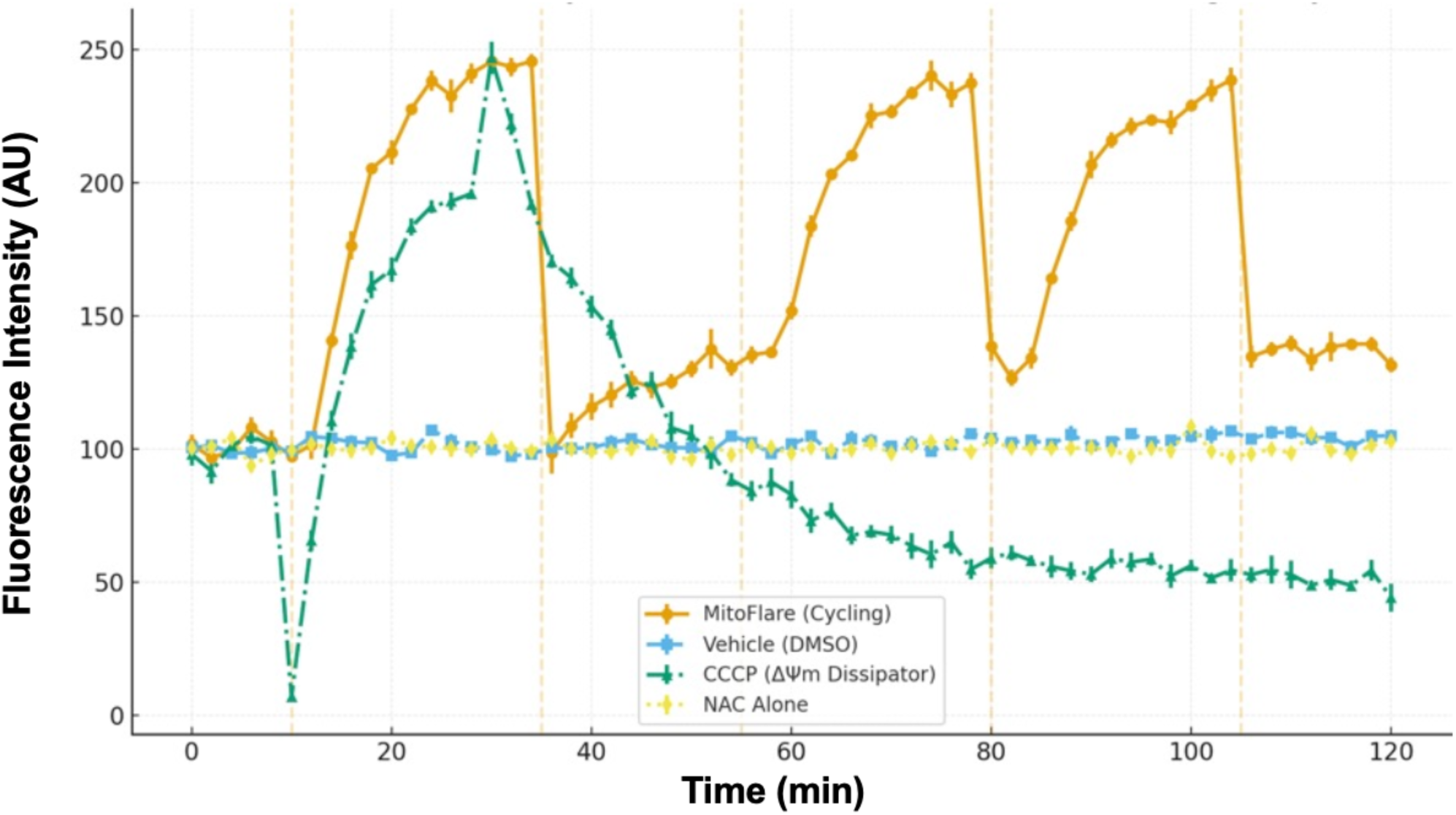
Cellular reversibility of MitoFlare in live PC12 cells. Kinetic fluorescence time course in MitoFlare-loaded PC12 cells (1 μM, 30 min) subjected to three cycles of mtROS induction (5 μM antimycin A at 10, 55, and 80 min) and quenching (0.5 mM NAC at 35, 80, and 105 min), showing ∼2.4-fold activation per cycle with 60–70% reversal and <8% cumulative signal loss over 120 min (orange; mean ± SEM, n=6 wells). Controls include vehicle (DMSO <0.1%; blue; <5% drift), CCCP (50 μM at 10 min; green; transient ∼2-fold surge followed by efflux-driven decay to ∼50% baseline), and NAC alone (0.5 mM at 10 min; yellow; no perturbation).

Control studies further validated the specificity of the observed cycling behavior. Vehicle-treated cells exhibited minimal drift (<5%), while NAC alone produced no measurable change in baseline fluorescence, indicating that reductive quenching requires prior ROS-mediated activation of the probe. In contrast, mitochondrial uncoupling with CCCP (10 μM) produced a transient ∼2-fold fluorescence increase, followed by an irreversible decline to ∼50% of baseline. This characteristic pattern is consistent with ΔΨm dissipation, subsequent loss of cationic probe retention, and collapse of localized mtROS signaling. These sharply divergent kinetic signatures—stable reversible cycling with antimycin A versus irreversible decay with CCCP—highlight the mechanistic fidelity and mitochondrial specificity of MitoFlare. These data demonstrate that MitoFlare undergoes rapid, repeatable oxidation–reduction transitions within living neurons, faithfully reporting mtROS dynamics without photobleaching, signal fatigue, or loss of mitochondrial localization. This reversible, mitochondria-retentive behavior establishes MitoFlare as a powerful tool for interrogating mitochondrial redox physiology and mtROS-driven mitophagy signaling in neuronal systems.

### 2.4. Quantitative Visualization of Mitophagy Using Dual MitoFlare–LysoTracker Live-Cell Imaging

To define the temporal architecture of mitophagy in PC12 cells exposed to metabolic stress or lysosomal dysfunction, we leveraged the unique capabilities of MitoFlare, a reversible and mtROS-responsive fluorogenic probe, to quantify three complementary metrics across a 350-minute live-cell imaging period. MitoFlare selectively illuminates mitochondria undergoing oxidative activation, enabling precise, non-genetic measurement of spatiotemporal organelle dynamics at each stage of the mitophagic cascade. The resulting dataset integrates the organelle proximity index (early mitochondrial recruitment toward lysosomes), Manders’ M1 coefficient (mid-stage mitochondrial–lysosomal fusion and content mixing), and the quenching/swelling index (late-stage lysosomal degradation). These measures reconstruct mitophagy flux with high temporal fidelity, as shown in Figure 4 (mean ± SEM, n = 4 biological replicates).

The stepwise progression of mitophagy is illustrated in Figure 3. MitoFlare–LysoTracker imaging captured heterogeneous yet ordered stages within the same field: red mitochondria enclosed within green lysosomes without yellow overlap (early engulfment prior to fusion); partial red–green colocalization with emerging yellow structures (active fusion and initiation of content mixing); and yellow-only autolysosomes lacking distinct red or green components (terminal digestion). These snapshots corroborate the temporal sequence quantified by the proximity index, Manders’ M1, and quenching/swelling metrics.

**Figure 3.**
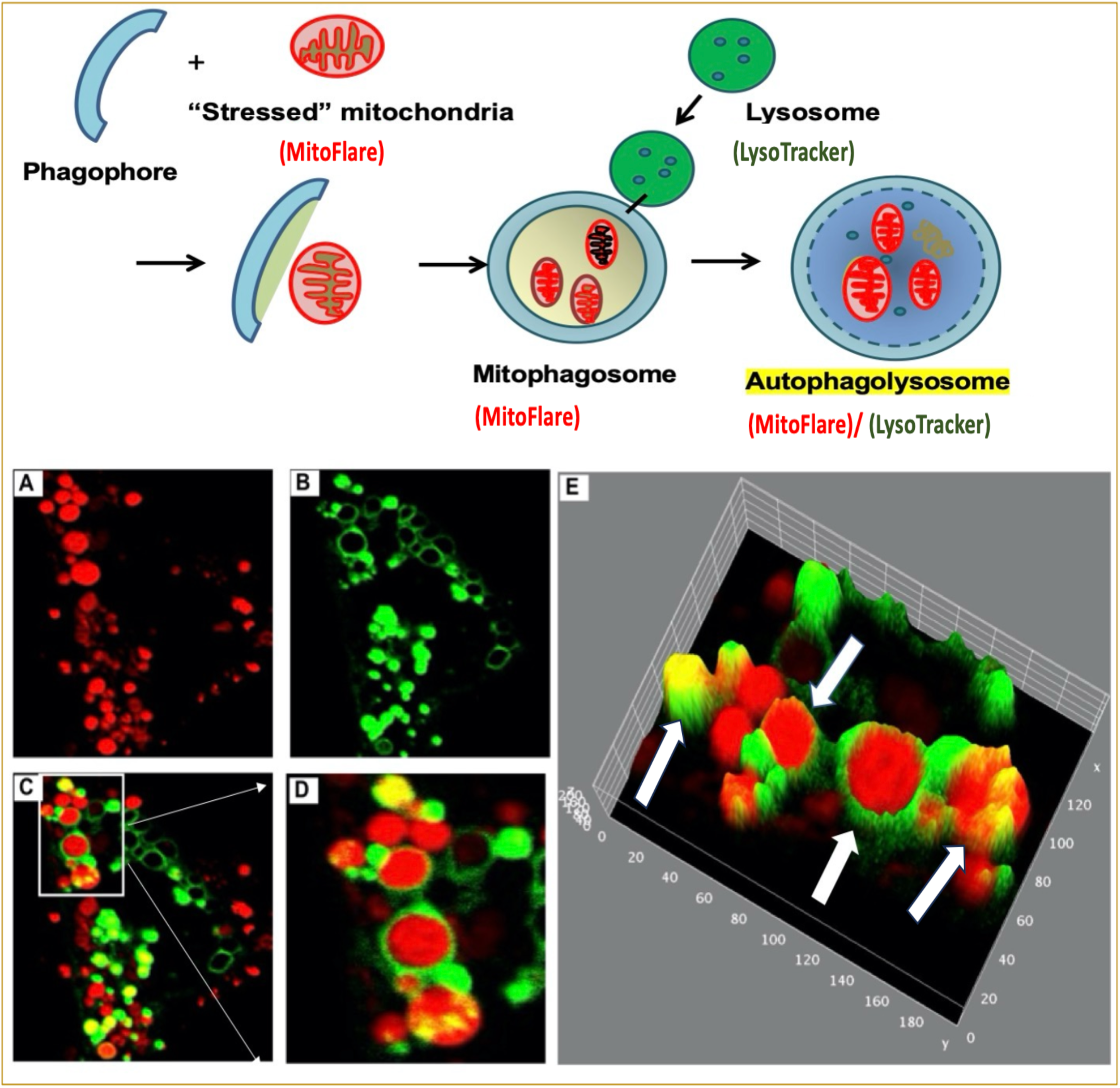
Confocal live-cell imaging of stepwise mitophagy progression visualized using MitoFlare and LysoTracker in PC12 cells under glucose deprivation. Representative confocal images demonstrate distinct stages of mitophagy within the same imaging field, reflecting asynchronous mitochondrial quality control among cells. In some cells, red mitochondria are trapped inside green lysosomes without yellow overlap, representing the early stage of mitophagy, where mitochondria are engulfed but not yet fused with lysosomes. Other cells exhibit red and green signals partially colocalized with emerging yellow fluorescence, corresponding to the active degradation phase, where mitochondrial and lysosomal compartments have fused and content mixing has begun. Several cells display yellow-only structures without distinct red or green components, indicative of the terminal mitophagy stage, where mitochondrial remnants have been fully degraded within mature autolysosomes. These snapshots capture the dynamic continuum of mitophagy from initial sequestration to complete mitochondrial digestion, validating MitoFlare’s capacity to resolve distinct temporal phases of mitochondrial turnover in live cells.

#### 2.4.1. Nutrient deprivation induces a fully phased and physiologically adaptive mitophagy program

Under nutrient deprivation, MitoFlare revealed a progressive and highly coordinated increase in the proximity index, reaching a maximum of approximately 0.65 during the 60–180 minute interval (Fig. 4A). This rise reflects the early stage of mitophagy—mitochondrial priming and recruitment—during which oxidatively stressed mitochondria undergo directed trafficking toward lysosomes. Because MitoFlare selectively illuminates mitochondria experiencing elevated mtROS, the increase in proximity indicates that ROS-positive mitochondria are selectively mobilized rather than all mitochondria indiscriminately. The decline after 180 minutes marks the resolution of organelle trafficking, consistent with the completion of early priming and the transition into later fusion and degradation steps.

**Figure 4.**
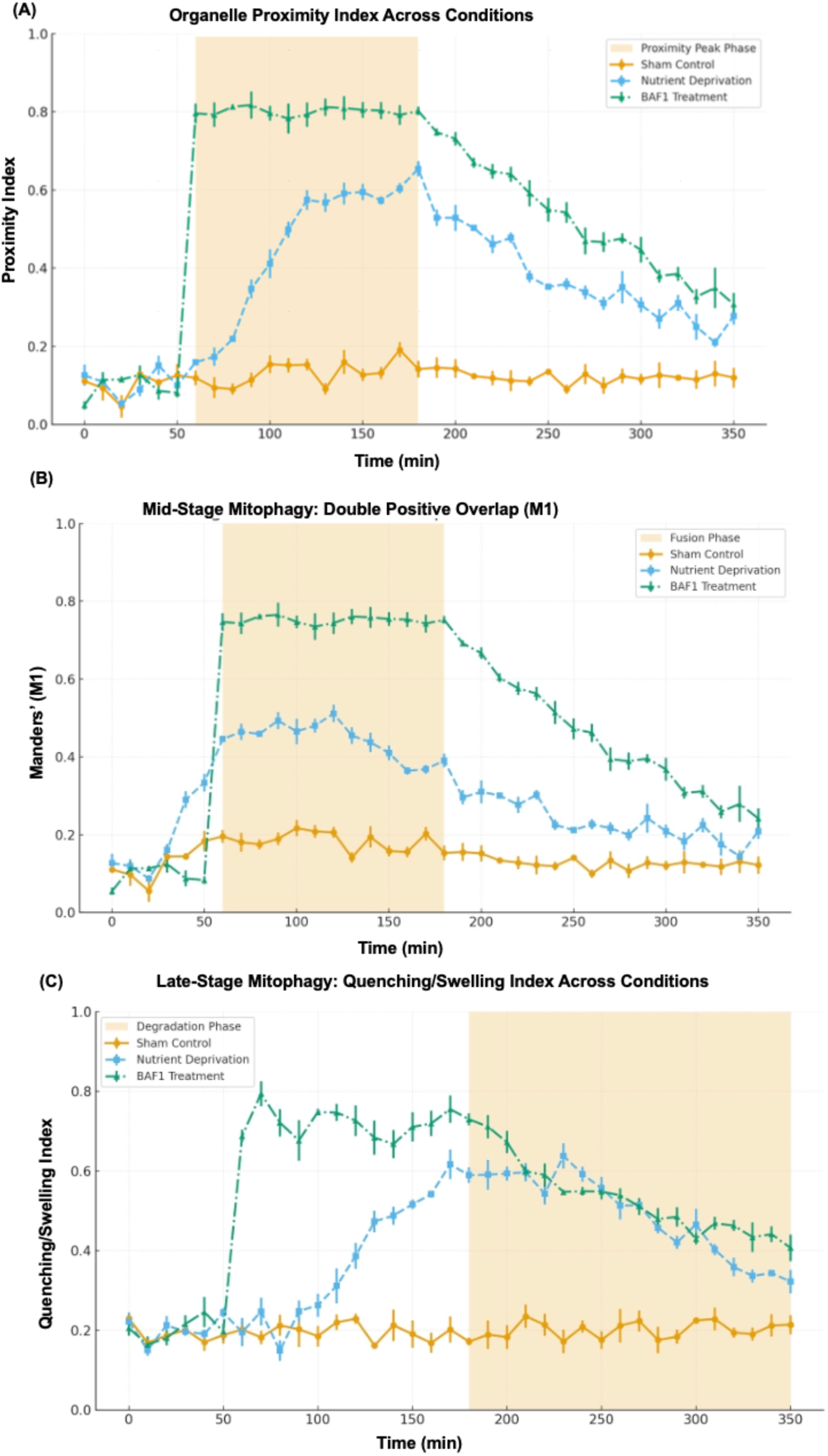
Quantitative temporal mapping of mitophagy progression using MitoFlare–LysoTracker dual imaging in live PC12 cells. (A) Early-stage mitochondrial–lysosomal engagement quantified by the Proximity Index. (B) Mid-stage fusion dynamics measured by Manders’ M1 coefficient (double-positive overlap). (C) Late-stage degradative processing assessed by the Quenching/Swelling Index. Across all panels, metrics were derived from MitoFlare–LysoTracker dual imaging (n = 120–160 cells per condition, four biological replicates). Each parameter captures a mechanistically distinct phase of mitophagy—early recruitment (A), fusion and content mixing (B), and terminal degradation (C)—providing a comprehensive temporal reconstruction of mitophagy dynamics in live neuronal cells.

Mid-stage fusion dynamics were captured through Manders’ M1 coefficient, which exhibited a well-defined increase followed by a gradual return toward baseline (Fig. 4B). This M1 rise reflects the formation of mitophagosomes—hybrid structures generated when autophagosomes and lysosomes dock and fuse. The pattern mirrors classical observations of LC3–LAMP1 juxtaposition during starvation-induced mitophagy in neurons, where PINK1–Parkin activity promotes ubiquitination of OMM substrates, thereby recruiting autophagic machinery for membrane tethering and fusion. As fusion proceeds, cargo enters the lysosomal lumen and moves toward degradation, which explains why M1 naturally declines as mitochondria are broken down and no longer generate a colocalized red–green signal.

The late-stage mitophagic phase was captured with high specificity by the quenching/swelling index (Fig. 4C). This metric reached a peak of approximately 0.60 at ∼170 minutes, reflecting the moment of maximal lysosomal engagement, when mitochondrial remnants first encounter an acidic and hydrolytically active lysosomal environment. The subsequent decline after 180 minutes corresponds to lysosomal acidification, cathepsin activation, and enzymatic digestion, processes that quench MitoFlare fluorescence due to oxidation of its reduced hydroxylamine state. Vesicular swelling at this stage results from osmotic changes accompanying macromolecular degradation, a phenotype frequently observed during autolysosome maturation. Thus, the decrease in signal reflects the biochemical hallmark of successful cargo breakdown. Together, proximity, fusion, and quenching signatures delineate a fully intact and physiologically adaptive mitophagy program progressing through coordinated phases of recruitment, fusion, and degradation.

#### 2.4.2. Lysosomal inhibition disrupts mitophagy completion despite amplified early recruitment

In contrast to nutrient deprivation, bafilomycin A1 (BAF1)—a selective v-ATPase inhibitor—disrupted the terminal stages of mitophagy and produced a fundamentally different kinetic signature. The proximity index rose rapidly and remained elevated (∼0.81) throughout the 60–180-minute window (Fig. 4A), demonstrating robust but unproductive recruitment of mitochondria to lysosomal compartments. This strong early rise reflects a compensatory response in which damaged mitochondria accumulate near lysosomes even though fusion and degradation cannot proceed. Mechanistically, the persistent proximity aligns with hyperactivation of RAB7 and SNARE docking pathways under lysosomal stress, a well-documented response in lysosomal storage disorders and Parkinsonian models, where impaired clearance triggers excessive trafficking to compensate for the downstream block.

Manders’ M1 coefficient reached a sustained plateau of approximately 0.77 and failed to return toward baseline until after 180 minutes (Fig. 4B). Importantly, M1 began to decrease only gradually beyond this point—a feature that reflects partial disengagement or remodeling of stalled hybrid organelles rather than true degradation. This behavior indicates that oxidatively stressed mitochondria indeed enter lysosome-associated compartments (yielding strong colocalization), but remain undegraded due to the absence of acidification. These long-lived “fusion-without-degradation” intermediates mirror the LC3- and LAMP1-positive autolysosome-like structures commonly observed in v-ATPase–blocked neurons.

The quenching/swelling index provided unambiguous confirmation of a terminal block in degradation. Values remained elevated (0.7∼0.8) long after the point when degradation normally occurs and declined only slightly to ∼0.41 over the full 350-minute period (Fig. 4C). This shallow decay reflects two key features: (i) failure of lysosomal acidification, which prevents MitoFlare quenching, and (ii) progressive accumulation of undegraded oxidative cargo, leading to swelling without breakdown. Because MitoFlare fluorescence persists in non-acidified compartments, the assay uniquely identifies this degradation failure. The high nadir (∼0.41) is characteristic of pathologically stalled mitophagic flux and parallels phenotypes reported in models of Parkinson’s disease, ALS, Niemann–Pick disease, and other lysosomal insufficiency syndromes.

#### 2.4.3. MitoFlare-enabled metrics reveal modular regulation of mitophagy

Across all three quantitative measures, BAF1 yielded a distinct pattern of mitophagy initiation in the absence of completion, while nutrient deprivation elicited a fully resolved mitophagic cycle. Statistical comparisons confirmed significant differences between stressed conditions and sham controls (proximity: P < 0.01; M1: P < 0.001; quenching/swelling nadir: P < 0.05 for nutrient deprivation and P < 0.001 for BAF1). Sham values remained unchanged across all time points, defining the homeostatic baseline for mitochondrial–lysosomal interactions.

This triad of metrics—made possible only by MitoFlare’s reversible mtROS-dependent activation—offers mechanistic granularity not attainable with irreversible fluorogenic dyes or genetically encoded reporters. The proximity index isolates early mitochondrial trafficking, Manders’ M1 resolves fusion and cargo incorporation, and the quenching/swelling index exclusively reports acidification-dependent degradation. Critically, these parameters demonstrate that early mitochondrial recruitment may appear normal even when terminal degradation is profoundly compromised, underscoring the necessity of multi-stage quantification to differentiate adaptive flux from pathological arrest.

#### 2.4.4. Mechanistic significance and translational implications

The staged dissociation between recruitment, fusion, and degradation phases highlights mitophagy as a modular process governed by discrete biochemical checkpoints. Nutrient deprivation accelerates mitochondrial priming and promotes efficient fusion and lysosomal degradation, representing an adaptive cytoprotective response to metabolic stress. In contrast, BAF1 selectively blocks the acidification checkpoint, yielding fused but undegraded organelles—a pathological signature shared across numerous neurodegenerative diseases, including Parkinson’s disease, ALS, and various lysosomal storage disorders.

MitoFlare’s reversible, redox-sensitive activation is essential for resolving these transitions, providing a dynamic, non-genetic, and high-fidelity readout of mitophagy progression that distinguishes early engagement from true flux completion. This capability is critical for evaluating new therapeutic strategies aimed at restoring lysosomal competence, enhancing mitochondrial turnover, or correcting flux defects in disease-relevant models.

## 3. Conclusion

This work establishes MitoFlare as a next-generation chemical reporter capable of resolving mitophagy with a level of temporal precision and mechanistic depth that has not been achievable with existing probes. By coupling reversible mtROS sensing to a mitochondria-targeted, long-wavelength fluorogenic scaffold, MitoFlare overcomes the inherent limitations of irreversible ROS dyes and the technical constraints of genetic autophagy reporters. When combined with LysoTracker, this platform enables continuous, non-genetic visualization of the complete mitophagic trajectory—from oxidative priming and organelle recruitment to mitochondria–lysosome fusion, to terminal acidification-dependent cargo degradation—within intact living cells.

Using this dual-probe system, we demonstrate that mitophagy unfolds through modular, quantifiable stages governed by discrete biochemical checkpoints. Nutrient deprivation activated a fully integrated and orderly mitophagy program, whereas lysosomal inhibition produced a signature of arrested flux characterized by persistent organelle tethering, exaggerated fusion intermediates, and a complete failure of degradative clearance. These results underscore the importance of measuring redox activation and lysosomal degradation in parallel, as early recruitment can remain deceptively intact despite profound downstream defects—an analytical blind spot of conventional autophagy markers.

Beyond methodological advancement, MitoFlare-enabled multiparametric imaging provides a conceptual and practical framework for dissecting mitochondrial quality-control pathways. Its reversible chemistry, high specificity, and non-genetic compatibility make it ideally suited not only for immortalized cell lines, but also for primary neurons, iPSC-derived cells, and ex vivo tissues where genetic manipulation is inefficient or perturbative. The ability to quantify mitophagic flux in real time positions this system as a powerful tool for identifying rate-limiting steps that fail in neurodegenerative, metabolic, and lysosomal storage diseases, and for evaluating compounds designed to restore mitochondrial turnover and cellular resilience.

In sum, MitoFlare establishes a new standard for chemical interrogation of mitophagy, enabling precise discrimination between adaptive mitochondrial clearance and pathological flux arrest. This platform offers broad utility for mechanistic discovery, therapeutic development, and high-content screening aimed at correcting mitochondrial dysfunction across diverse disease contexts.

## 4. Limitation

While this work establishes MitoFlare as a highly responsive and reversible probe for resolving mitochondrial redox dynamics and mitophagy progression, several opportunities remain to further extend its applicability. The current study focuses on PC12 neuronal cells, a widely used and well-characterized model that enables precise control of mitochondrial stress and live-cell cycling. Expanding future experiments to additional neuronal subtypes and primary cells will allow exploration of how MitoFlare performs across diverse mitochondrial phenotypes and physiological environments, thereby broadening its utility.

The solution-phase and cellular cycling assays presented here demonstrate robust reversibility and excellent retention within polarized mitochondria. As MitoFlare is applied to increasingly complex tissue systems, it will be valuable to refine calibration strategies that relate fluorescence changes to absolute redox parameters, enabling deeper quantitative comparisons across biological models. Such advances are readily achievable using complementary tools such as redox clamping, metabolic perturbation profiling, and genetically encoded sensors.

The imaging-derived metrics developed here—including proximity, Manders’ overlap, and late-stage degradation indices—provide a powerful framework for parsing mitophagy stages in live cells. Future studies employing higher-resolution imaging modalities or automated segmentation pipelines may further enhance spatial precision and accelerate throughput, positioning these metrics for broad adoption in screening and mechanistic studies.

Finally, although MitoFlare performs robustly under the conditions tested, ongoing work aimed at examining its behavior under prolonged imaging or under extreme mitochondrial depolarization will help define its boundaries and enable optimizations tailored for specific applications such as long-term neuronal tracking or in vivo imaging.

## 5. Materials and Methods

### 5.1. Reagents and General Procedures

All reagents were purchased from Sigma-Aldrich (St. Louis, MO, USA) unless otherwise stated. Phosphate-buffered saline (PBS, pH 7.4), hydrogen peroxide (H₂O₂, 30% w/v), ascorbic acid, and catalase (Aspergillus niger, ≥2,000 U/mg) were used without further purification. Ultrapure water (18.2 MΩ·cm) was obtained from a Milli-Q system. Spectroscopic measurements were performed in 1-mL quartz cuvettes (10-mm path length).

### 5.2. Synthesis of MitoFlare

#### 5.2.1 Rhodamine B piperazine-azido-PEG₄ amide (Compound 4)

4: To a stirring solution of Rhodamine B piperazine amide (538 mg, 1.08 mmol), N_3_-PEG_4_-COOH (291 mg, 1 mmol), and Et3N (327 mg, 3.24 mmol) in anhydrous CH_2_Cl_2_ (30 mL) was added HBTU (409 mg, 1.08 mmol). The mixture was stirred for 3 h at room temperature and diluted with CH_2_Cl_2_ (100 mL). The CH_2_Cl_2_ layer was washed with brine (30 mL × 2). The organic phase was dried over anhydrous Na_2_SO_4_, filtered, and concentrated. Flash column chromatography of the residue on silica gel (CH_2_Cl_2_:MeOH, 20:1) afforded the compound 4 as a purple foam (310 mg, 59.1%). ^1^H NMR (400 MHz, CDCl_3_) δ ppm 7.64-7.52 (m, 2H), 7.58-7.49 (m, 1H), 7.35-7.23 (m, 1H), 7.29-7.19 (m, 1H), 7.04-6.79 (m, 2H), 6.84-6.72 (m, 2H), 3.71 (d, *J* = 10.6 Hz, 2H), 3.67-3.47 (m, 20H), 3.46-3.22 (m, 10H), 2.68-2.49 (m, 4H), 1.28 (t, *J* = 7.1 Hz, 12H). ^13^C NMR (100 MHz, CDCl_3_) δ ppm 178.45, 167.93, 157.89, 157.85, 155.86, 135.21, 132.30, 130.35, 128.51, 114.00, 112.52, 96.23, 87.97, 73.05, 70.49, 70.34, 70.02, 67.04, 53.85, 50.79, 48.32, 46.26, 45.79, 41.91, 41.23, 34.15, 33.19, 13.87, 12.75. HRMS (FAB+) *m/z* 828.4608, calcd for C_48_H_59_N_8_O_5_ 828.0471.

Rhodamine B piperazine-PEG₄-triazolyl-N,N-bis(propargyl)methanaminium (Compound 5): To a solution of compound 4 (323 mg, 0.39 mmol) and tripropargylamine (153 mg, 1.17 mmol) in anhydrous acetonitrile (20 mL) was added copper(I) iodide (21 mg, 0.11 mmol). The mixture was stirred for 3 h at room temperature, concentrated under reduced pressure, and diluted with CH_2_Cl_2_ (50 mL). The CH_2_Cl_2_ layer was washed with brine (10 mL × 2). The organic phase was dried over anhydrous Na_2_SO_4_, filtered, and concentrated. Flash column chromatography of the residue on silica gel (CH_2_Cl_2_:MeOH, 20:1) afforded the desired compound 5 as a purple foam (109 mg, 30.5%).^1^H NMR (400 MHz, CDCl_3_) δ ppm 7.67 (s, 1H), 7.71-7.59 (m, 2H), 7.60-7.47 (m, 1H), 7.35-7.24 (m, 1H), 7.15 (d, *J* = 9.5 Hz, 2H), 6.97-6.78 (m, 2H), 6.77-6.66 (m, 2H), 4.47 (t, *J* = 7.4 Hz, 2H), 3.80 (t, *J* = 4.8 Hz, 2H), 3.75 (s, 2H), 3.74-3.65 (m, 2H), 3.61-3.44 (m, 20H), 3.42-3.37 (m, 8H), 3.34-3.20 (m, 4H), 2.62-2.45 (m, 2H), 2.21 (t, *J* = 2.4 Hz, 2H), 1.23 (t, *J* = 7.1 Hz, 12H). ^13^C NMR (100 MHz, CDCl_3_) δ ppm 170.58, 167.85, 157.86, 155.83, 144.18, 135.23, 132.20, 130.24, 124.46, 114.54, 113.93, 96.25, 78.83, 73.70, 70.35, 69.64, 67.03, 50.26, 48.12, 46.23, 42.04, 33.24, 12.73. HRMS (FAB+) *m/z* 915.5133, calcd for C_52_H_67_N_8_O_7_ 916.1534.

Synthesis of MitoFlare (Compound 6): A solution of Rhodamine B piperazine–PEG₄–triazolyl–N,N-bis(propargyl)methanaminium (compound 5, 165 mg, 0.18 mmol) and 4-azido-TEMPO (108 mg, 0.55 mmol) in anhydrous acetonitrile (10 mL) was treated with copper(I) iodide (11 mg, 0.06 mmol) under an inert atmosphere. The reaction mixture was stirred at room temperature for 3 h, concentrated under reduced pressure, and diluted with CH₂Cl₂ (50 mL). The organic phase was washed twice with brine (2 × 10 mL), dried over anhydrous Na₂SO₄, filtered, and evaporated to dryness. The crude residue was purified by flash column chromatography on silica gel (CH₂Cl₂/MeOH = 20:1) to yield compound 6 (MitoFlare) as a purple foam (87 mg, 36.8%). ^1^H NMR (400 MHz, CDCl3) δ ppm 8.15 (brs), 7.75 (br.s), 7.36 (br.s), 7.23 (br.s), 6.84 (br.s), 4.68 (br.s), 3.71 (br.s), 2.72 (br.s), 1.46 (br.s). ^13^C NMR (100 MHz, CDCl_3_): No peaks. MS (FAB+) m/z 1310.9, calcd for C_70_H_10_1N_16_O_9_ 1309.79. The ^1^H NMR spectrum exhibited broad singlets within a narrow chemical shift range, lacking clear multiplicities or integrations—a behavior characteristic of molecules containing proximate nitroxide radicals. In this case, the two TEMPO moieties in compound 6 (C₇₀H₁₀₁N₁₆O₉⁺) induce strong paramagnetic relaxation effects, resulting in extensive line broadening and coalescence of distinct proton resonances. The absence of observable ^13^C NMR signals is consistent with the presence of unpaired electrons, which markedly shorten T₂ relaxation times and suppress carbon signal detection in conventional ^13^C NMR, particularly in CDCl₃ without specialized low-temperature or relaxation-optimized techniques. The FAB⁺ mass spectrum (m/z 1310.9; calcd 1309.79) agrees closely with the theoretical monoisotopic mass of the cationic species, confirming the molecular composition and structural integrity of MitoFlare.

### 5.3. Solution-Phase Redox Cycling Assay

#### 5.3.1. Fluorescence Cycling Protocol

To assess intrinsic reversibility of MitoFlare, a temperature-controlled iterative redox assay was conducted. A 1 mM DMSO stock (<0.1% DMSO final) was diluted to 10 μM in PBS immediately before use. Following 5-min equilibration at 25°C, continuous baseline fluorescence was recorded (λex = 560 nm; λem = 588 nm; 5-nm slit widths).

Sequential oxidation–reduction cycles were initiated by micropipette addition of reagents with gentle mixing. For Cycle 1, H₂O₂ was added at t = 5 min (final 100 μM) and the signal was monitored for 10 min until plateau. Ascorbic acid (final 1 mM) was then added at t = 15 min to quench the probe, followed by 10 min monitoring. The procedure was repeated for Cycles 2 and 3 (H₂O₂ at t = 25 and 45 min; ascorbate at t = 35 and 55 min), yielding a 60-min time course sampled every minute (n = 3 replicates). Controls included (i) MitoFlare-only vehicle (PBS/DMSO) and (ii) catalase (10 U/mL added at t = 10 min) to verify enzymatic H₂O₂ degradation and switching specificity.

#### 5.3.2. Spectral Data Analysis

Fluorescence traces were exported via FluorEssence (v3.7) and processed in GraphPad Prism (v10.0). Curves were baseline-corrected using pre-cycle means and smoothed by LOESS regression (span = 0.1). Quantitative indices were defined as:

Activation fold-change per cycle:

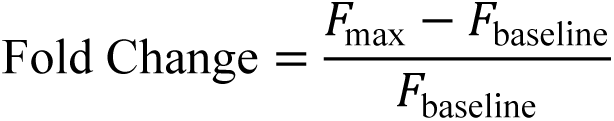

Quenching efficiency:

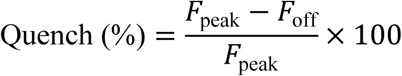

Cumulative attenuation across cycles:

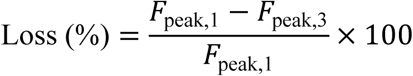

One-way ANOVA with Tukey’s post hoc test (α = 0.05) was applied to compare fold-changes. Normality (Shapiro–Wilk) and homoscedasticity (Bartlett’s test) assumptions were met.

### 5.4. Electron Paramagnetic Resonance (EPR) Validation

Nitroxide cycling was corroborated by electron paramagnetic resonance. Spectra of post-cycle reaction mixtures (100 μL, equivalent to 100 μM MitoFlare) were recorded at 25°C on a Bruker EMX X-band spectrometer (9.8 GHz) using a 1 G modulation amplitude, 20 mW microwave power, and 100 kHz modulation frequency. Samples were loaded into quartz capillaries (50 μL). Double-integrated EPR intensities were normalized to a 100 μM TEMPO standard. Redox recovery was calculated as:

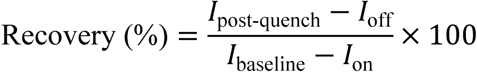

### 5.5. Cellular ROS Induction–Quenching Assay

#### 5.5.1. PC12 Cell Culture and Differentiation

PC12 cells were cultured in RPMI-1640 supplemented with 10% horse serum, 5% fetal bovine serum, and 1% penicillin–streptomycin at 37°C, 5% CO₂. For differentiation, cells were seeded at 5 × 10^4^ cells/well in phenol red–free RPMI-1640 on collagen IV–coated black optical-bottom 96-well plates and treated with 100 ng/mL nerve growth factor (NGF) for 5–7 days. Neuronal differentiation was confirmed by neurite extension >50 μm in >80% of cells.

#### 5.5.2. Probe Loading and Colocalization Markers

Differentiated cells were equilibrated for 15 min at 37°C in phenol red–free RPMI. MitoFlare was added to a final concentration of 1 μM (1 mM DMSO stock; <0.1% DMSO) and incubated for 30 min. Excess dye was removed by two gentle washes with 100 μL medium. For lysosomal colocalization, 200 nM LysoTracker Green was included during the final 15 min of loading.

#### 5.5.3. Live-Cell Kinetic Fluorimetry

Mitochondrial ROS was induced by adding antimycin A at 10, 55, and 80 min (final 5 μM). The resulting “on” phase was monitored for 25 min. Reductive quenching was triggered by NAC at 35, 80, and 105 min (final 0.5 mM), followed by 20-min monitoring. Between cycles, wells were washed twice and replenished with fresh medium. Three full cycles were completed over ∼120 min (n = 6 wells/condition). Cell-free wells provided background correction.

#### 5.5.4. Control Conditions

Control experiments were performed to validate the specificity of the observed fluorescence changes. Vehicle-treated cells (DMSO <0.1%) were used to assess photophysical stability and exhibited less than 5% signal drift over the course of the assay. Treatment with NAC alone at 0.5 mM tested the integrity of the off state and did not alter basal MitoFlare fluorescence, indicating that quenching requires prior ROS-dependent activation. In contrast, exposure to 10 μM CCCP was used to evaluate ΔΨm dependence and produced a transient approximately twofold increase in fluorescence followed by an irreversible decline to about 50% of baseline, consistent with mitochondrial depolarization and loss of cationic probe retention.

#### 5.5.5. Cellular Data Analysis

Quantitative metrics included:

Activation fold-change:

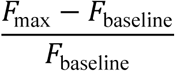

Quenching efficiency:

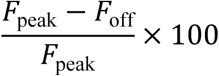

Cumulative attenuation across cycles:

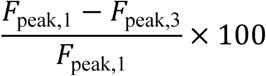

Replicate means ± SEM were graphed. Inter-cycle differences were assessed via one-way ANOVA with Dunnett’s post hoc test (α = 0.05). Shapiro–Wilk and Levene’s tests confirmed normality and homogeneity of variance. MitoFlare–LysoTracker/MitoTracker colocalization was quantified using Fiji (v2.3.0; Coloc2). Cell viability exceeded 90% by MTT assay.

#### 5.5.6. Quantitative Metrics for Mitophagy Flux

Mitophagy progression was quantified from time-lapse confocal imaging using three orthogonal parameters:

##### (i) Proximity Index — Early Priming

A unitless metric (0–1) reflecting pre-fusion recruitment of mtROS-positive mitochondria toward lysosomes:

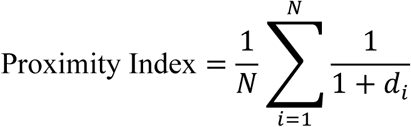

where dᵢ is the Euclidean distance (μm, capped at 5 μm) between each MitoFlare-positive punctum and its nearest LysoTracker-positive boundary. Values >0.4 indicate priming (Δindex/Δt > 0.01/min).

##### (ii) Manders’ M1 — Mid-Stage Fusion

Manders’ coefficient quantifies the fraction of red (mitochondrial ROS) signal overlapping green (lysosomal) ROIs:

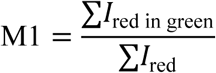

Threshold-independent analyses (Coloc2) were used. M1 > 0.5 indicates robust fusion; mean latency from priming >0.4 to M1 > 0.5 was ∼45 min.

##### (iii) Quenching/Swelling Index — Late-Stage Degradation

A composite index integrating probe quenching and lysosomal expansion:

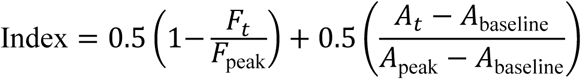

where Fₜ is MitoFlare intensity and Aₜ lysosomal area. Efficient turnover corresponds to nadir values <0.3.

Integrated Flux Score:

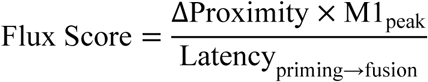

yielding a dimensionless measure (0.3–0.7 range) of mitophagy efficiency. Field-level values (10–15 fields/dish) were analyzed by mixed-effects ANOVA (lme4, R).

## Author Contributions

The manuscript was written through contributions of all authors. All authors have given approval to the final version of the manuscript. ‡These authors contributed equally.

## ACKNOWLEDGMENT

We sincerely thank the NIH (R01HL163159, Z.S.), NIH (R15 EB035866, L.B), and the American Heart Association (AHA, grant #1807047, LB), GLRC-ICC (L.B., R01805) for their generous financial support. We also extend our great gratitude to Dr. Rick Koubek for his encouragement and unwavering support throughout our project.

